# Fast in vivo ^23^Na imaging and T_2_^*^ mapping using accelerated 2D-FID magnetic resonance spectroscopic imaging at 3 T: Proof of concept and reliability study

**DOI:** 10.1101/2020.05.25.114918

**Authors:** Ahmad A. Alhulail, Pingyu Xia, Xin Shen, Miranda Nichols, Srijyotsna Volety, Nicholas Farley, M. Albert Thomas, Armin M Nagel, Ulrike Dydak, Uzay E. Emir

## Abstract

**Purpose:** To implement an accelerated MR-acquisition method allowing to map sodium T_2_^*^ relaxation and absolute concentration within skeletal muscles at 3T.

**Methods:** A fast-2D density-weighted concentric-ring-trajectory ^23^Na-MRSI technique was used to acquire 64 time-points of FID with a spectral bandwidth of 312.5 Hz from a 2.5 x 2.5 mm^2^ in-plane resolution within about 15 minutes. The fast relaxing ^23^Na signal was localized with a single-shot, inversion-recovery based, non-echo (SIRENE) OVS method. The sequence was verified using simulation and phantom studies before implementing it in human calf muscles. Within two same-day sessions, 2D-SIRENE-MRSI (UTE = 0.55 ms) and 3D-MRI (UTE = 0.3 ms) data were acquired. The T_2_^*^ values were fitted voxel-by-voxel using a bi-exponential model for the 2D-MRSI data. Within-subject coefficients of variation were estimated for both acquisition methods.

**Results:** The MRSI-FID data allowed for fast and slow T_2_^*^ mapping of the calf muscles in vivo with minimal sensitivity reduction. The spatial-distributions of ^23^Na concentration for both in vivo MRSI and 3D-MRI acquisitions were significantly correlated (r = 0.7, *P*<0.001). The test-retest results rendered high reliability for both MRSI (CV = 5%) and 3D MRI (CV = 6%). The mean 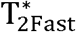 in calf muscles was 0.7 ± 0.1 (contribution fraction = 37%), while 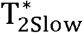 was 13.2 ± 0.2 ms (63%). The mean absolute muscle ^23^Na concentration calculated from the T_2_^*^-corrected data was 28.6 ± 3.3 mM.

**Conclusion:** The proposed MRSI technique is a reliable technique to map sodium’s absolute concentration and T_2_^*^ within a clinically acceptable scan time at 3T.

## 1 INTRODUCTION

Sodium (^23^Na) plays a crucial role in preserving many vital functions in our bodies. Under healthy conditions, ^23^Na concentration stays within certain ranges but varies among tissue types.^1^ The increase in these ranges is a sign of health disorder or physiological changes. In skeletal muscles, the rise in ^23^Na concentration was used as a biochemical marker for several diseases such as hypertension^2^, diabetes^3^, or muscular channelopathies.^4,5^ It was also tested for monitoring early therapeutic responses.^6^ The physiological changes have also been observed by using ^23^Na-MR experiments to assess the influence of exercise. An increase in muscle sodium concentration has been noticed after the exercises.^7^ In a similar study, this has been suggested to be a result of an increase in the transverse relaxation time (T_2_), which eventually led to an increase in the detected MR signal.^8^

^23^Na possesses 3/2 nuclear spin and its relaxation is determined by the quadrupolar interaction (QI = the interaction between the ^23^Na nuclear quadrupole moment and the local electric field gradients), and the containing environment.^9,10^ In an aqueous solution, the time-averaged QI equals zero, and the spins move freely and tumble very fast. Subsequently, ^23^Na exhibits a relatively long monoexponential T_2_. However, within a biological system, ^23^Na spins interact with the surrounding macromolecules, and a time averaged QI ≠ 0 is possible. As the mobility of the ^23^Na gets more restricted, the transverse relaxation becomes biexponential.^10^ The monoexponential relaxation decay within in-vitro aqueous ^23^Na solutions was estimated to be ranging between 20-50 ms.^11,12^ In-vivo, a monoexponential ^23^Na relaxation of 55-65 ms was also measured within the CSF.^1,13^ In human muscles, a fast transverse relaxation time 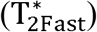 ranges between 0.5–3.0 ms, and slow transverse relaxation time 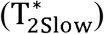 between 12–28 ms were observed.^14^ This variation in relaxation modes and magnitude between the in-vivo and in-vitro environments can cause a T_2_ quantification bias if no correction is performed. Similarly, the longitudinal relaxation time (T_1_) of ^23^Na is also short. At 3T, the T_1_ of skeletal muscles has been estimated to be about 29 ms.^4^ Thus, a careful relaxation measurement is necessary to conduct accurate quantification studies and can also yield useful physiological information.

Within the few conducted in-vivo skeletal muscle studies, the used ^23^Na-relaxometry techniques suffer either from a limitation in their spatial precision (large voxel covering different tissue types) or from their impractical acquisition time. The intact skeletal muscles T_2_^*^ values have been measured previously using a non-localized FID method.^8,15^ Such relaxometry methods estimate the averaged value from a large scanned area, which may include different tissues in addition to the muscles. Moreover, because of this partial volume effect, the estimated value may not be enough to perform a voxel-wise relaxation correction. To achieve the relaxometry study with more spatial precision, a 3D-UTE acquisition technique was used with images acquired at different echo times, which consumes a long scan time (9 min per image), and with a low in-plane spatial resolution (6 x 6 mm^2^)^7^. Alternatively, a multi-echo GRE sequence has been used to measure the T_2_^*^ over the entire slice of interest with higher resolution and shorter total time (~14 minutes).^16^ However, since GRE starts with a slice-selective gradient, the minimum possible first echo was 1.9 ms, which may not be short enough to detect the fast decaying component of the T_2_^*^. To allow shorter TEs and avoid the lengthy slice-selective gradients, outer volume suppression (OVS) methods such as single-shot, inversion-recovery based, non-echo (SIRENE)^17^ and FIDLOVS^18^ have been proposed. However, these methods have not been applied and tested for ^23^Na imaging.

Therefore, the goal of this work was to develop an accelerated method with an early acquisition start to estimate in vivo T_2_^*^ values of muscle tissues in a voxel-wise manner at 3T while maintaining high spatial and temporal resolution. To attain this, we are proposing an accelerated density-weighted concentric ring trajectory (DW-CRT) MRSI acquisition to measure in vivo ^23^Na relaxation times in the lower leg muscles with a high sampling frequency. Additionally, to mitigate the long TE limitation when using slice-selective gradients, the SIRENE method^17^ was used instead to reach UTE. Before conducting an in-vivo repeatability study, the localization method was tested using simulation and in-vitro experiments.

## 2 METHODS

### 2.1 Sequence design

The sequence begins with a pair of slice-selective gradients and adiabatic full passage inversion pulses (hyperbolic secant (HS20) pulses, B_1_ peak of 19.2 μT, a pulse duration of 12.8 ms, and a thickness of 100 mm) applied to invert the magnetization bands outside the desired slice along the z direction and followed by spoiler gradients. After a delay chosen to null the inverted magnetization (TI = ln[2] x tissue’s T_1_, 20 ms here), pairs of wide-bandwidth adiabatic half passage OVS saturation pulses (HS20, B_1_ peak of 29 μT, a pulse duration of 2.56 ms, and thickness of 100 mm) and gradients along z-axis were applied. The net effect is to eliminate the magnetization within the bands at each side of the slice of interest (SOI) along the z direction. The FID is then measured from the remaining magnetization within the SOI after a square excitation pulse of 240 μs duration (see Figure 1). To accelerate data collection, the k-space data is collected by using a fast density-weighted concentric ring trajectory (DW-CRT) acquisition similar to that implemented in Chiew et al.^19^

**FIGURE 1.**
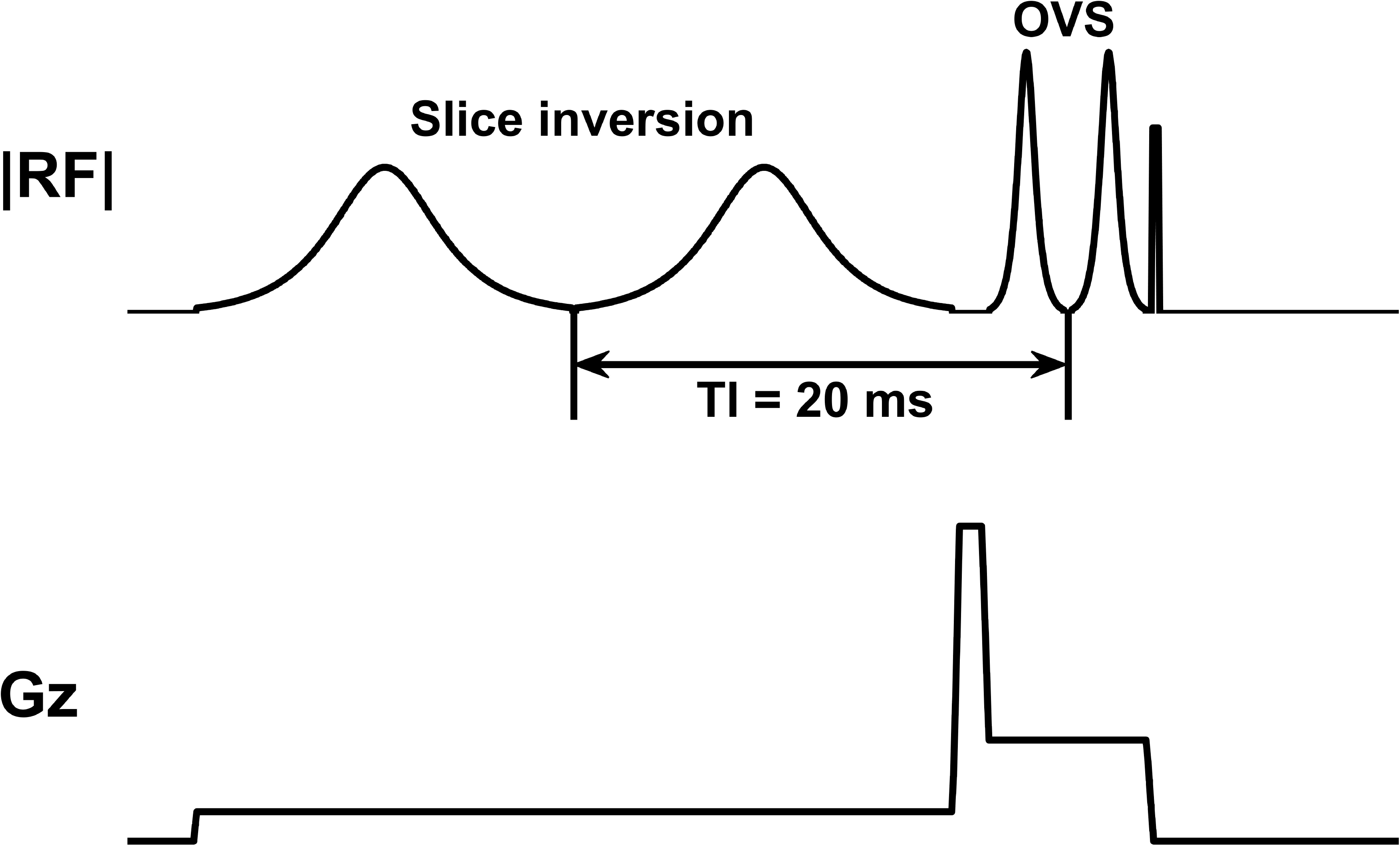
Pre-readout MRSI pulse sequence. To suppress the unwanted region outside the slice of interest (SOI), two OVS bands are assigned before the FID excitation. These OVS bands along the z-direction are applied after two selective 180° inversion recovery pulses covering the area outside the SOI at inversion time (TI) = ln(2) x 29 ms = 20 ms, where 29 ms is the muscles T1.^4^ Directly after the OVS band pulses, a nonselective 90° excitation pulse is applied before starting the FID-MRSI readout

### 2.2 Outer volume suppression bands performance evaluation

#### 2.2.1 Simulation

To evaluate the OVS bands’ performance in eliminating the entire ^23^Na signal outside the SOI, the OVS pulse was simulated on SpinBench (HeartVista, Inc. Menlo Park, CA). Along with the approximated skeletal muscle sodium NMR properties at 3T (gyromagnetic ratio = 11.25 MHz/T, T_1_ = 29 ms, fast T_2_^*^ = 0.5 ms, and slow T_2_^*^ = 12 ms), the identical MRSI experimental parameters were used for the simulation software. The resulted thickness, sharpness, and residual signal amplitude over the generated spatial profile were used to judge and optimize the bands’ parameters.

#### 2.2.2 Phantom experiment

To test the optimized OVS bands parameters on the scanner, a phantom study was conducted. Four phantoms (bottles) were prepared with different known ^23^Na concentrations (10, 20, 30, 40 mM), which will also serve a reference for the in vivo signal calibration. To mimic the in-vivo T_1_ of sodium, 2.9 g/L CuSO_4_ was added to each phantom.

As demonstrated in Figure 2A, to ensure that no signal is coming from outside the SOI, a scan was acquired while two ^23^Na phantoms of the highest concentration (30 and 40 mM) were placed outside the SOI, within the OVS bands. The remaining phantoms of lower concentration (10 and 20 mM) were placed in the center of the SOI. For comparison, ^23^Na images with a 3D-MRI sequence scan covering the same SOI thickness were also acquired.

**FIGURE 2.**
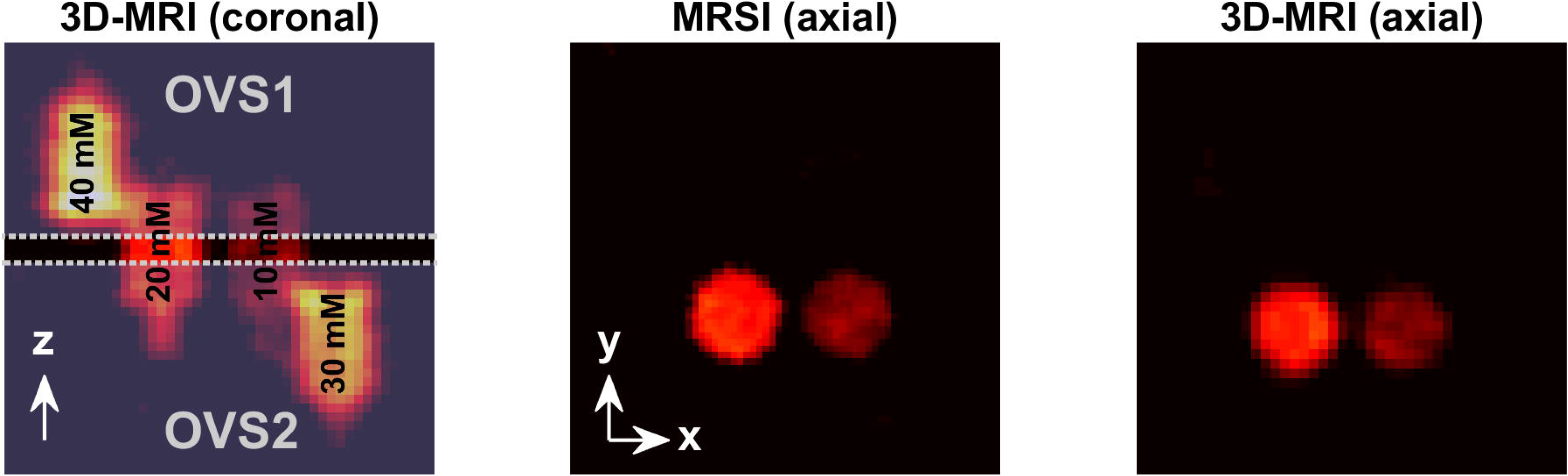
Phantom evaluation of the OVS localization. Two ^23^Na phantoms (10 and 20 mM) were placed at the center of the slice of interest (SOI), which were located between the 2 OVS bands. Two additional phantoms of higher concentrations (30 and 40 mM) were placed outside the SOI, within the OVS bands and away from the center. As shown in the axial images, the MRSI signal is obtained only from the phantoms within the SOI, resulting in an image that is very similar to that produced by the 3D-MRI sequence

### 2.3 In vivo experiment

#### 2.3.1 Human subjects

In vivo calf muscle scans were performed in four subjects [1 male and 3 females; age 22-40 years (median = 26 years); body mass index (BMI) = 24 ± 3 kg/m^2^]. The study was conducted in accordance with the institutional review board of Purdue University. Before being scanned, an informed written consent was obtained from all the subjects.

#### 2.3.2 Repeatability study

In order to evaluate the reliability of the proposed MRSI method, test-retest scans were performed. The subjects were asked to lie in a head-first supine position with their left leg within a ^23^Na coil. The maximum circumference of their lower leg was positioned to be centered in the middle of the coil. Additionally, to make sure that both scans were acquired from the same slice, an ink marker was used to draw a line on the leg region (~10.5-11.5 cm below the knee joint, based on the subject) where the scanner laser was centered for the first scan. After a short break (about 5 minutes) outside of the scanner, the repeat scan was acquired using the same scanning protocol.

#### 2.3.3 Scanning parameters

The data were collected using a 3T Siemens Prisma MR system (Siemens Healthineers, Germany) and a frequency-tuned mono-resonant ^23^Na-transmit/receive birdcage knee coil (32.6 MHz, Stark-Contrast, Erlangen, Germany).

The ^23^Na data were measured through the default shim currents of the MR system (“Tune Up” under shim settings).

For the ^23^Na-MRSI study, inversion pulses with TI of 20 ms were applied, followed by the two OVS pulses on the z-direction preceded by the excitation RF pulse. The FID DW-CRT MRSI was implemented with an alpha of 1 (to improve SNR)^19^ and the following parameters: matrix size = 96 x 96, field of view (FOV) = 240 x 240 mm^2^, resolution = 2.5 x 2.5 x 20 mm^3^ (nominal resolution = 0.125 mL), flip angle (FA) = 90°, acquisition delay (time from the center of excitation pulse to the first FID time point) = 0.55 ms, repetition time (TR) = 650 ms, temporal samples = 64, number of rings = 48, points-per-ring = 64, spectral bandwidth = 312.5 Hz, spatial interleaves = 4, and readout duration = 204.8 ms. The scan performed with 8 averages, which resulted in a total acquisition time (TA) of 15 minutes and 24 seconds.

For comparison, ^23^Na-MRI was collected with a density-adapted 3D radial acquisition sequence^20^ with UTE = 0.3 ms, TR = 100 ms, FA = 90°, 1 average, 4 x 4 x 4 32 mm^3^ nominal resolution, and FOV = 256 x 256 mm^2^, and TA = 6.66 min. Five axial slices were averaged to cover the same MRSI slice. These 3D MRI images will serve as a reference to assess the spatial distribution of the MRSI maps.

To get an anatomical image suitable for segmentation, ^1^H images were acquired using the integrated body coil with a T_1_-FLASH sequence of TR/TE = 250 ms/2.46 ms, FA = 60°, 2 averages, 0.6 x 0.6 x 10 mm^3^ resolution, and FOV = 200 x 200 mm^2^.

All the above sequences were planned to collect data from the same axial slice placed at the scanner isocenter. Additionally, shimming using the ^1^H coil was done before the sodium measurements.

#### 2.3.4 Post-processing

The reconstruction of the MRSI data were performed in MATLAB (MathWorks, Natick, MA, USA). The gridding and the Fast Fourier Transform were done using the Nonuniform FFT (NUFFT) method.^21^ In addition to the DW-CRT trajectory^19^, a Hanning filter was applied in k-space for density compensation.

The B_0_ inhomogeneity was corrected by calculating the ^23^Na ΔB_0_ maps as described in Gast et al.^22^ Here, the ^23^Na ΔB_0_ maps were calculated based on the first two ^23^Na-MRSI phase-unwrapped images (*θ*_*TE*_2_,*unwrapped*_ and *θ*_*TE*_1_,*unwrapped*_) acquired at TE_1_ = 0.55 ms, and TE_2_ = 3.75 ms, as follows:

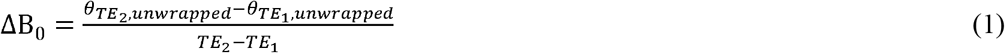

These ΔB_0_ maps were used to reconstruct field-corrected ^23^Na-MRSI images using a fast iterative image reconstruction method.^23^ Finally, we applied low rank approximations for spatial-spectral filtering of reconstructed ^23^Na MRSI data.^24^

The 3D-MRI data were reconstructed using a MATLAB tool designed for the radial sequence. To reconstruct the 3D-MRI magnitude images, the k-space data were density compensated before being re-gridded with an oversampling ratio of two using a Kaiser-Bessel kernel,^25^ and Fourier transformed by a conventional fast FFT. The data were filtered with a Hanning filter.

#### 2.3.5 Fitting of T_2_^*^

The acquired FID data from the leg region were fitted to a biexponential decay to calculate the 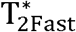 and 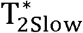 relaxation time components:

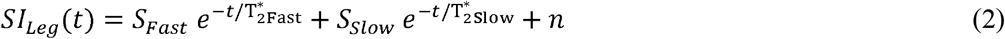

Here, *S_Fast_* and *S_Slow_* are the contributions to the initial signal from the fast and slow components, respectively, *t* indicates the FID points collection time, and *n* represents the offset level.

The initial (undecayed) signal equals the sum of *S_Fast_* and *S_Slow_*. Thus, their contribution fractions were represented as:

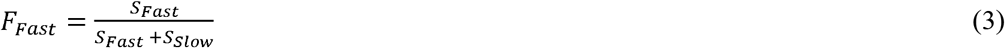

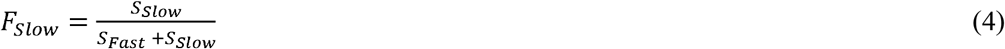

For the reference bottles, *Ref*, the FID curves were fitted to a monoexponential decay:

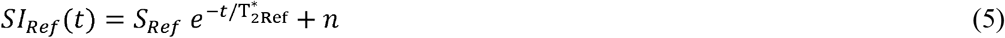

#### 2.3.6 Quantification

The ^23^Na concentration maps in mM were reconstructed by calibrating the signals of the reference bottles to their corresponding known concentrations. The resulting signal-to-concentration linear equation was used to calibrate the signals within the leg and to estimate the concentration in mM.

#### 2.3.7 Muscle segmentation

To assess the sodium concentration and T_2_^*^ values within the human calf muscles, the high-resolution T1-weighted image was used to manually draw regions of interest (ROIs) over each of the seven main large muscles (Figure 3). The borders of these ROIs were determined by tracing the boundaries of their corresponding muscle. Following, The ROIs were down-sampled and co-registered to each sodium map to evaluate the ^23^Na spatial distribution. Voxels close to the main blood vessels were avoided. The subcutaneous fat region was segmented similarly.

**FIGURE 3.**
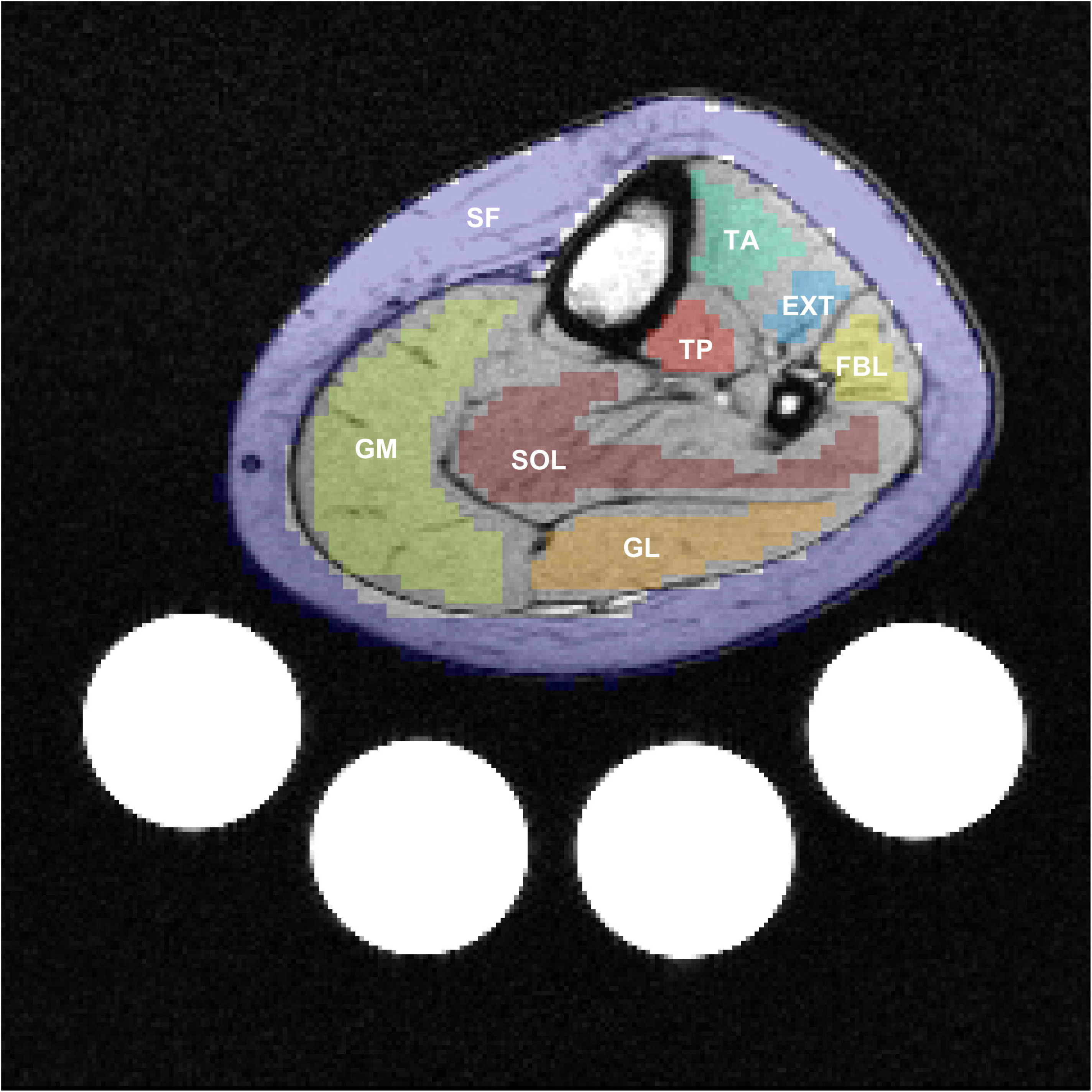
Representative segmented regions of interest (ROIs). Calf muscle and subcutaneous fat (SF) ROIs were drawn based on their high-resolution T1 axial ^1^H-image, which allows clear anatomical features for segmentation. Seven ROIs were drawn to cover the main human calf muscles: SOL, Soleus; FBL, Fibularis; EXT, Extensor longus; TA, Tibialis anterior; GM, Gastrocnemius medialis; GL, Gastrocnemius lateralis; and TP, Tibialis posterior muscles. The shown ROIs are presented in the MRSI resolution and overlaid over their corresponding high-resolution T1 anatomical image

#### 2.3.8 Statistical analysis

The Spearman regression analysis was performed to assess the signal spatial distribution agreement between the two used acquisition techniques within the segmented ROIs. The MRSI signals were extrapolated to their estimated values at 0.3 ms to match the TE of 3D-MRI).

To evaluate the repeatability of the MRSI and 3D-MRI methods, coefficient of variance (CV), intraclass correlation coefficient (ICC), and Bland–Altman analyses were utilized.

### 3 RESULTS

#### 3.1 Outer volume suppression bands performance evaluation

The result of the simulation (Figure 4) showed that using a TI of 20 ms within a region of muscle provides a sharp suppression profile. The spatial profile in Figure 4 represents the ^23^Na magnetization after the 90° excitation. While the signal within the OVS bands was totally suppressed, the magnetization within the SOI has full transverse magnitude and resulted in a slice thickness of 20 mm.

**FIGURE 4.**
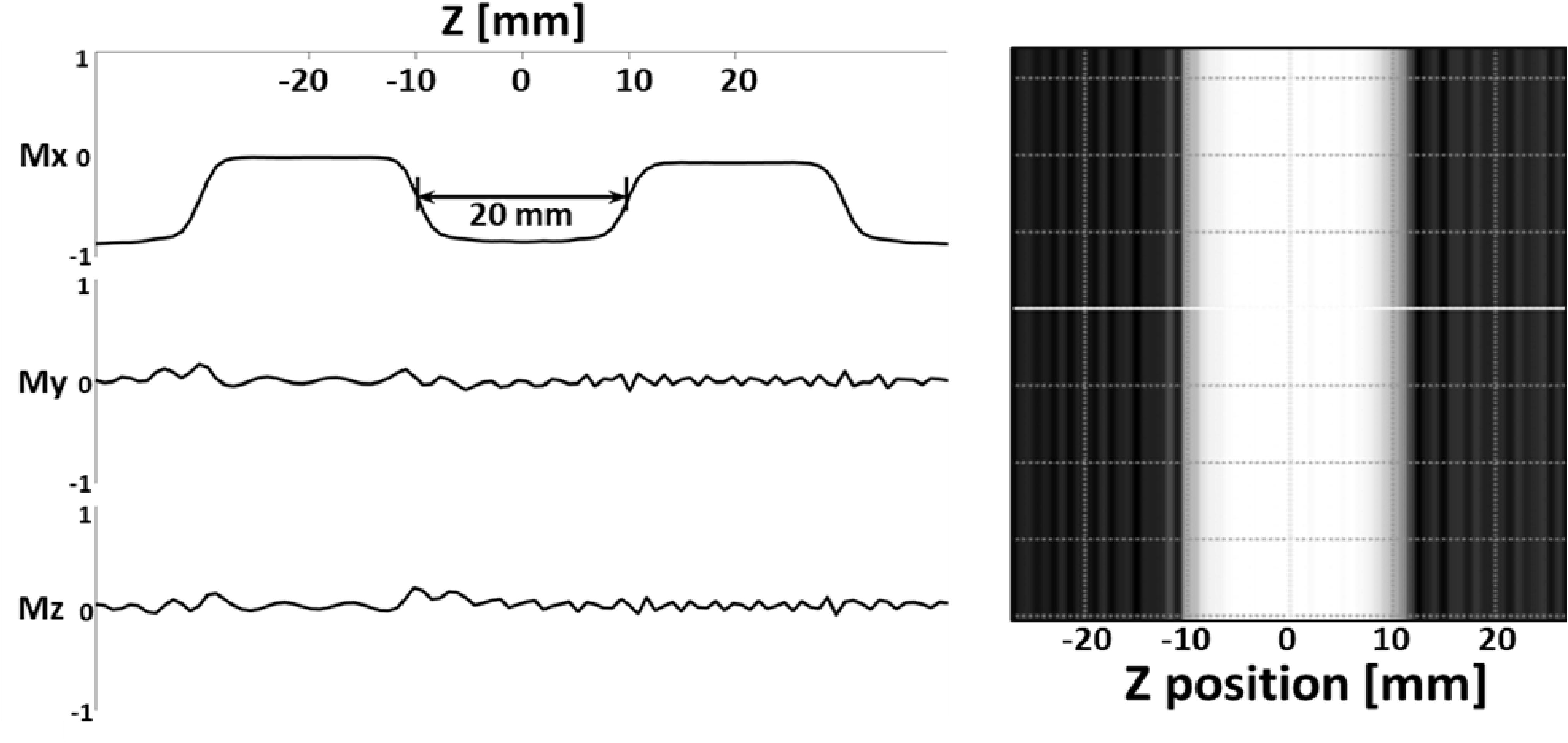
Simulated data shows a sharp spatial profile along the z-direction when using an inversion time of 20 ms

In-line with the simulation, the phantom study demonstrated that the signals from the high ^23^Na concentration phantoms (30 and 40 mM) placed within the OVS bands were totally suppressed (Figure 2B). In contrast, the signals of the low concentration phantoms (10 and 20 mM) within the SOI did not have any contamination and were in agreement with the 3D-MRI sequence (Figure 2C).

#### 3.2 In vivo experiment

As shown in Figure 5, comparison of the ^23^Na concentration spatial distribution between the MRSI and the 3D MRI (data from 64 muscle and subcutaneous fat ROIs from all subjects) resulted in a regression line of slope = 1.01 that is close to the unity line with a correlation coefficient (r) = 0.7 (*P* < 0.001). Additionally, the Bland–Altman analysis comparing between MRSI and the 3D-MRI showed a bias of 0.7 mM with a CV of 9 % (Figure 5D).

**FIGURE 5.**
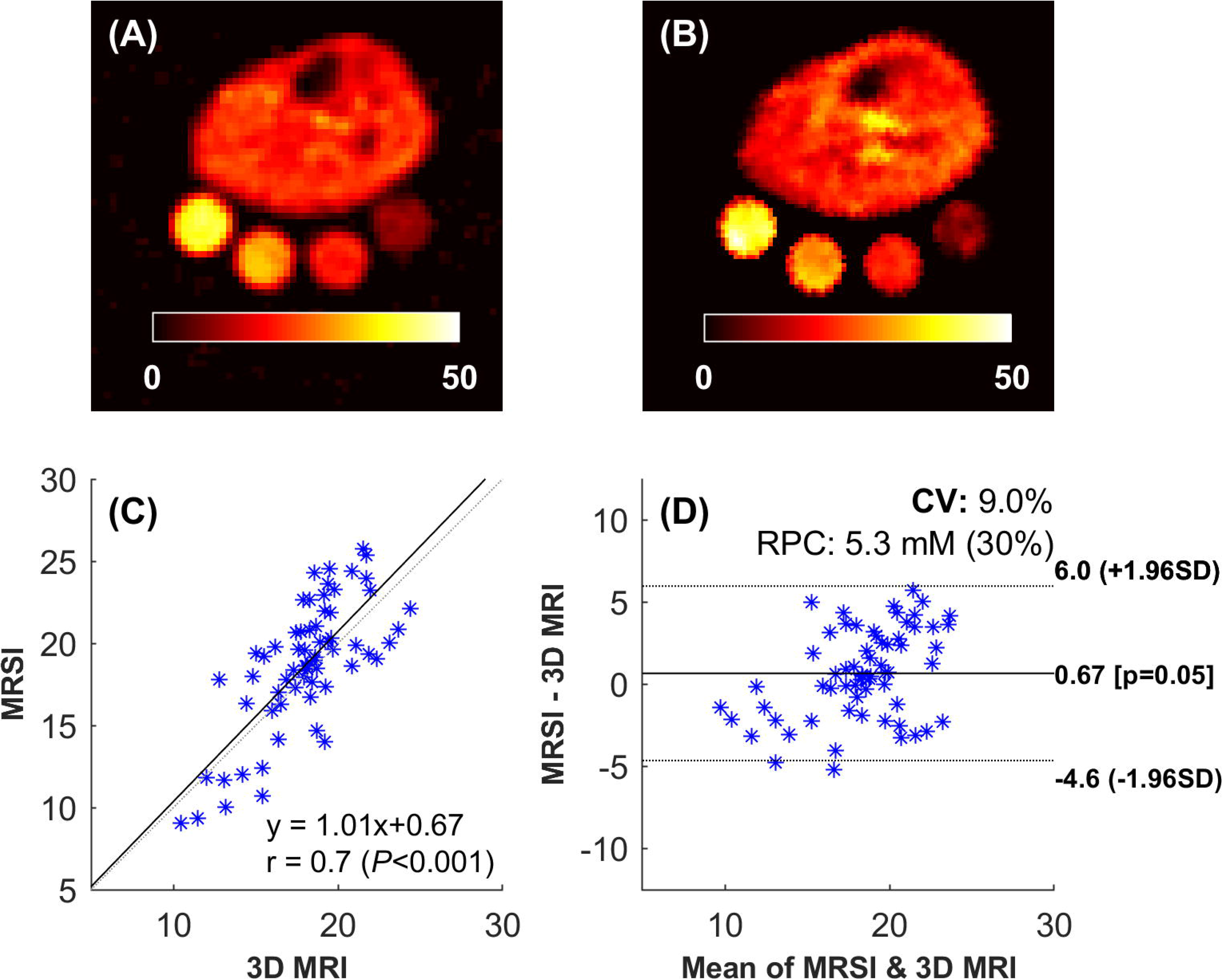
Correlation of MRSI with 3D MRI. A, A 3D ^23^Na-MRI map example. B, A ^23^Na-MRSI map example for the same subject (the signal was extrapolated to represent data at 0.3 ms, which is the 3D-MRI TE). C, Results of the regression analysis comparing the normalized mean signal (normalized to the phantom signal) of the MRSI and the 3D MRI within 64 regions of interest from all subjects data (7 muscles and one subcutaneous fat ROIs x 4 subjects x 2 scans). D, Bland-Altman analysis comparing the MRSI and the 3D-MRI results

The results of the biexponential T_2_^*^ values (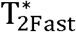 and 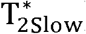) and their fractions are summarized in Table 1, which also includes the T_2_^*^ corrected concentration within each ROI. Fast and slow T_2_^*^-maps and their corresponding signal fraction maps are illustrated in Figure 6. The T_2_^*^ correction was demonstrated by showing sodium maps before and after correction with their difference (Figure 7A-C). Additionally, Figure 7D shows an example of two signals with similar initial MR signals diverging with time due to their different transverse relaxation in different environments. The ^23^Na spins decay monoexponentially within the reference phantom (blue) and biexponentially within the muscle (green).

**TABLE 1.**
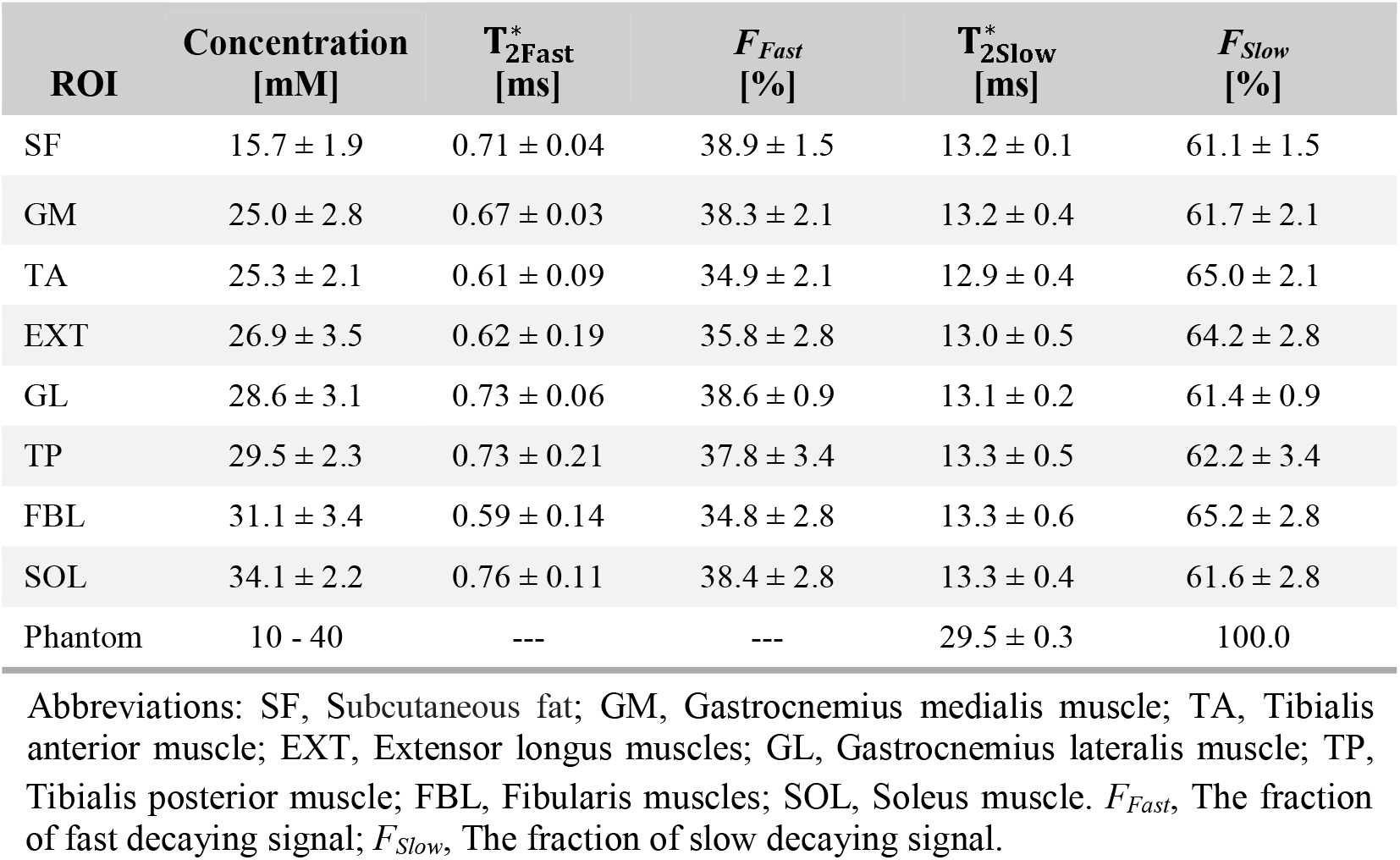
Regions of interest absolute ^23^Na concentrations, T_2_^*^ values, and signal fractions

**FIGURE 6.**
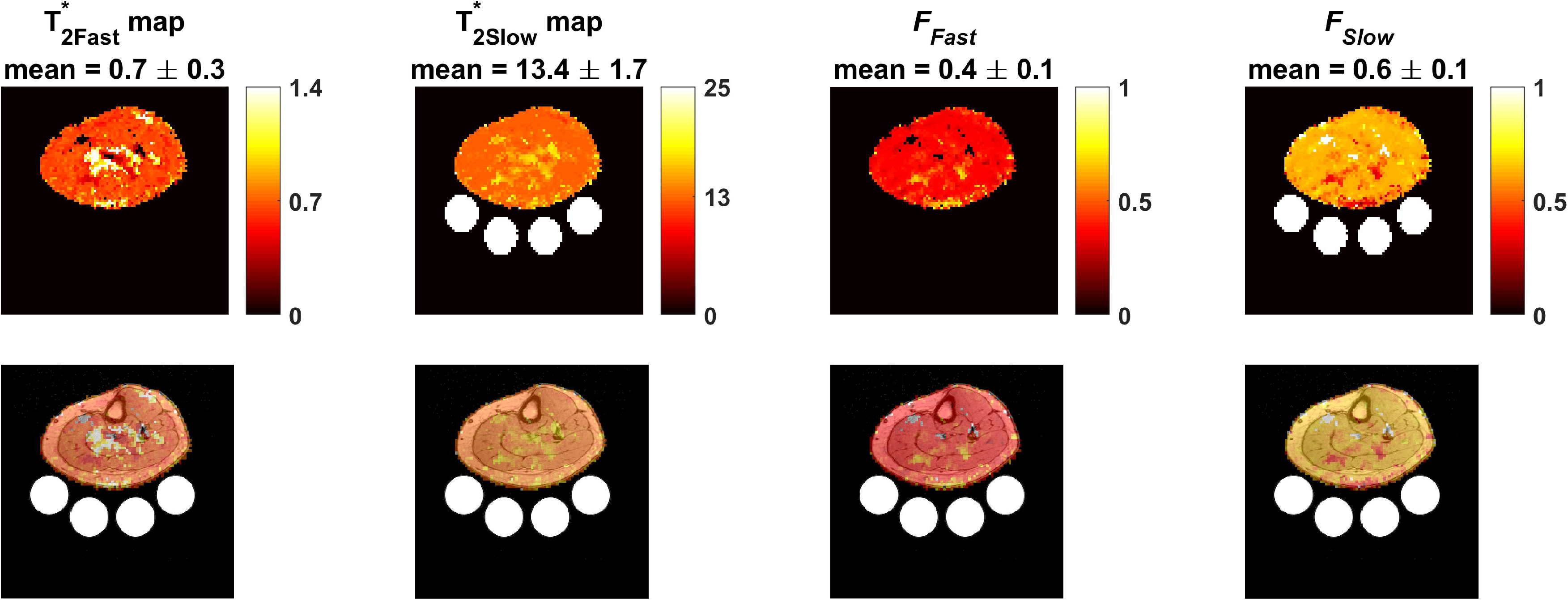
Representative relaxation maps. Top panel: fast and slow T_2_^*^ maps, and their corresponding signal fraction (*F_Fast_*, and *F_Slow_*). Their mean values from the entire leg slice (without the bottles) are listed above their maps. Bottom panel: the same maps overlaid on their anatomical images

**FIGURE 7.**
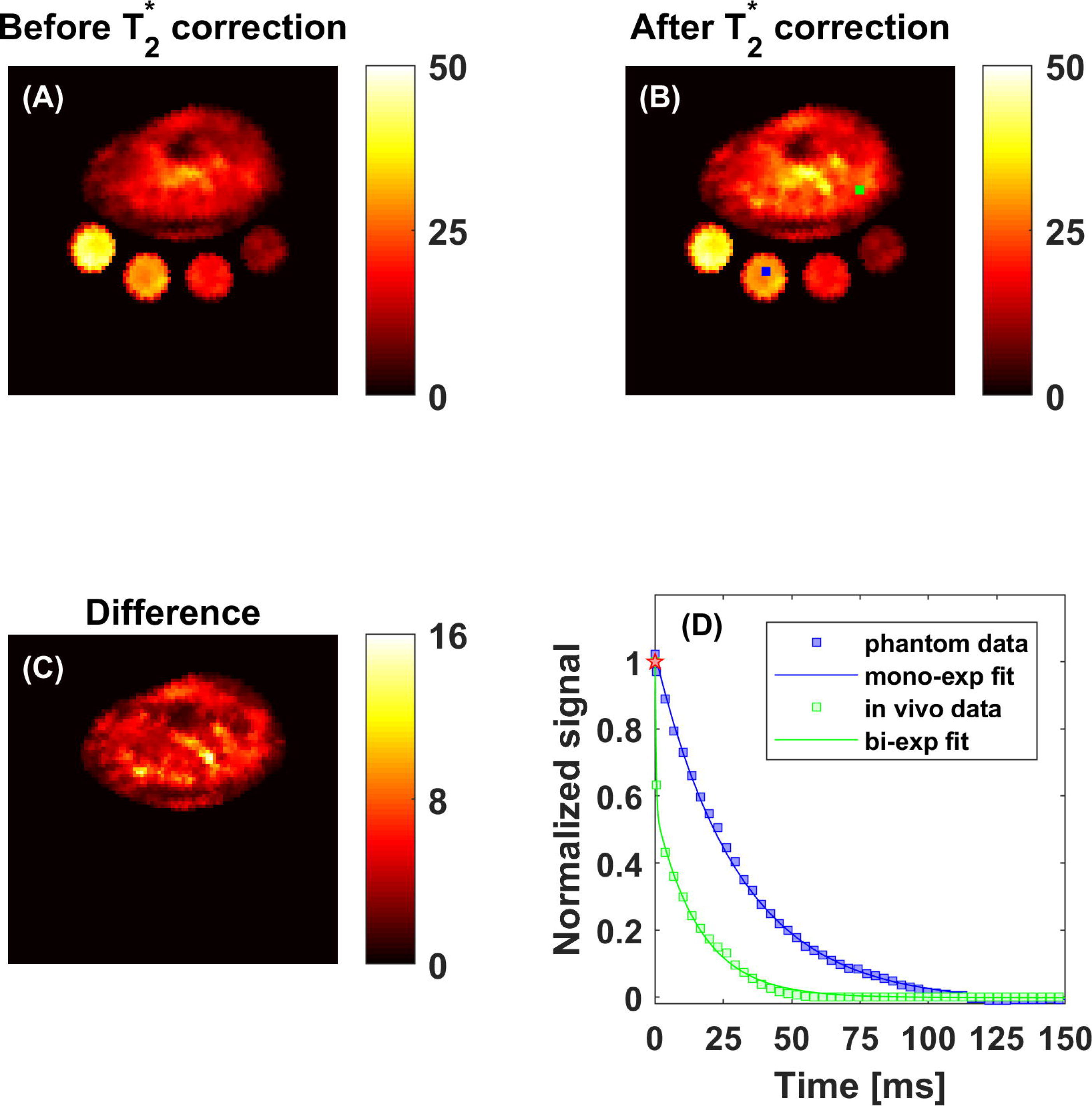
Illustrations of the importance of the relaxation correction. The difference between the data acquired at 0.55 ms (A) and the data after correction (B), shows a large improvement in concentration estimation. In the difference map (C), about 5 mM difference in muscles was found. The red star (D) represents the proton density signal corresponding to 30 mM absolute concentration. The fitting example of leg voxel (green box, B and D) and quantification reference voxel (blue box, B and D) with this absolute concentration shows how their signal can diverge with time before fully decaying

A representative set of baseline and repeated scan maps of the MRSI ^23^Na concentration (before and after T_2_^*^ correction) and their corresponding 3D-MRI maps are provided in Figure 8. The repeatability results of the muscle and subcutaneous fat ROIs from all subjects are shown in Figure 9. The CV was calculated as 4.2, 5.2, and 5.9 %, and the ICC as 0.98, 0.95, and 0.89 for the MRSI data before correction, after correction, and the 3D MRI data, respectively.

**FIGURE 8.**
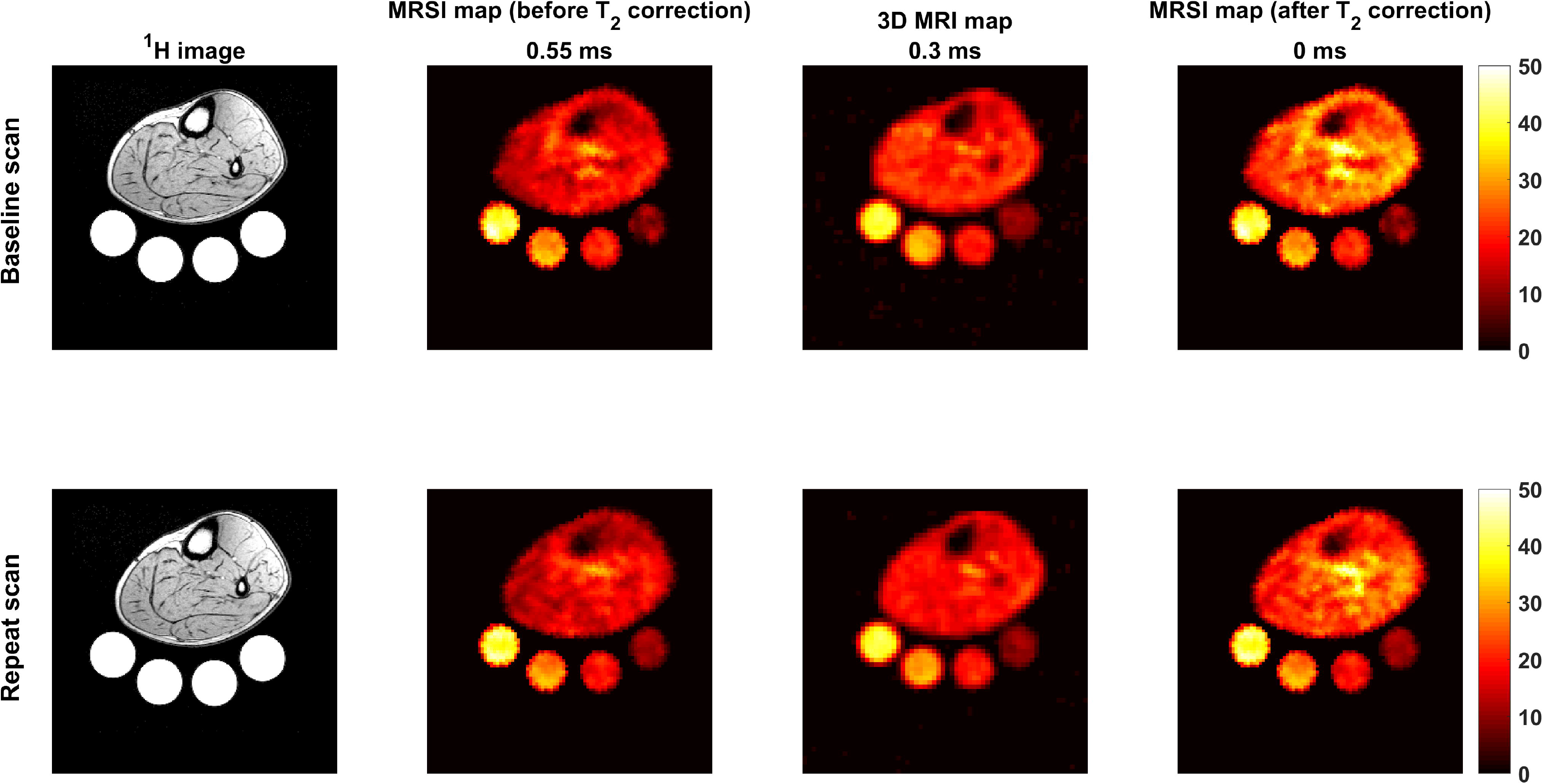
Data example of baseline and repeat scans

**FIGURE 9.**
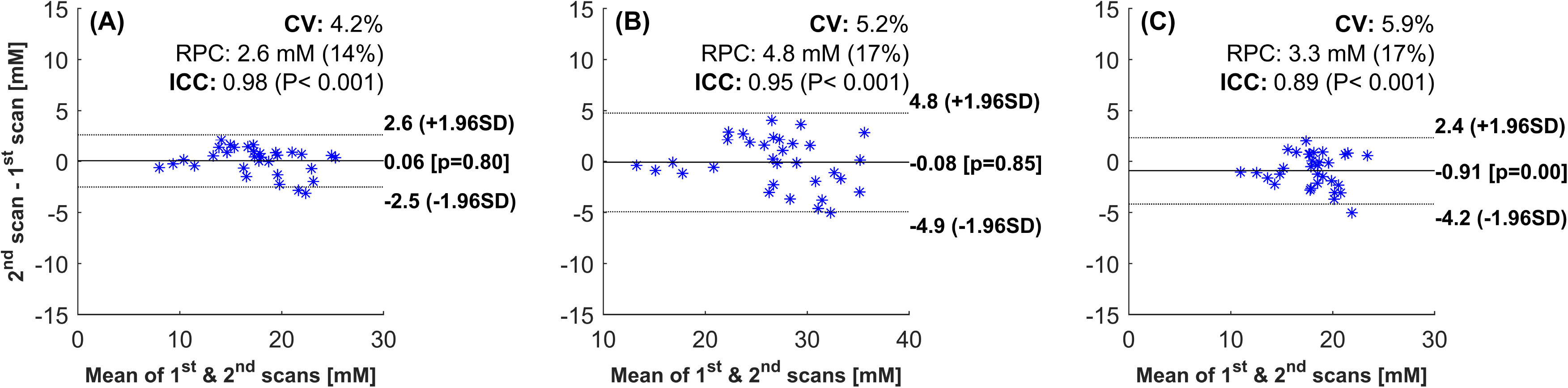
Evaluation of repeatability by Bland-Altman analysis. A, The results from the MRSI data before T_2_^*^ correction. B, The results from the MRSI data after the T_2_^*^ correction. C, The 3D-MRI data results. The graphs represent the variability of the measured data (muscles and subcutaneous fat ROIs from all subjects, 7 muscles and one subcutaneous fat ROIs x 4 subjects) between 1^st^ and 2^nd^ scans. The coefficient of variation (CV), reproducibility coefficient (RPC), systematic bias, and intraclass correlation coefficient (ICC) values are listed on each plot

## 4 DISCUSSION

In this work, we have introduced a novel method to acquire ^23^Na 2D-FID data with short acquisition delay (0.55 ms) with a good quality comparable to a well-established 3D acquisition method. The proposed method provides additional time points in the FID to calculate the T_2_^*^ and correct for its effects.

To reduce the delay time while performing a 2D-measurement, we utilized a single-shot, inversion-recovery based, non-echo (SIRENE) method based on OVS bands^17^. In this study, we implemented the technique with a novel accelerated k-space trajectory, DW-CRT,^19^ and showed its feasibility for the ^23^Na acquisition. The simulation and phantom studies showed that the proposed technique could be used as an alternative to the slice-selective gradient approach in which achieving an ultra-short TE is difficult.

Considering the variation in the parameters of the MRSI and the 3D-MRI sequences, the spatial distribution of their signal was in a good agreement (Figure 5C) with a small bias of 0.67 mM that was found with a 9 % CV. This variability between the two sequences results is expected due to their different spatial resolution (MRSI: 2.5 x 2.5 mm^2^; 3D-MRI: 4 x 4 mm^2^), and spatial response function. ^19,20^ Moreover, the field inhomogeneitiy correction was only performed on the MRSI reconstruction. A recent study with a similar 3D sequence, but with double echo to measure B_0_ and correct for its inhomogeneity at 3T, showed a small enhancement in the results.^26^

In this work, we provided an accelerated technique to conduct voxel-wise T_2_^*^ measurements within an acceptable scan time (15 minutes) while maintaining the spatial quality. It is worth mentioning that recently a number of studies have shown the feasibility of voxel-wise T_2_^*^ mapping in the brain using UTE 3D acquisition techniques.^27–31^ However, these techniques were conducted with fewer time points (8-38 echoes), and require long acquisition times (26 min - 1 h). Moreover, many of these methods were either done with lower spatial resolution, at a higher magnetic field, or done with both. If our proposed MRSI method was applied under these conditions, further reduction in scan time can be achieved.

The estimated mean muscle relaxation times (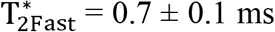 of *F_Fast_* = 37 ± 2 %; 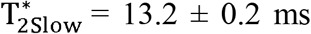 of *F_slow_* = 63 ± 2 %) are in line with previously reported values measured using single voxel ^23^Na MRS (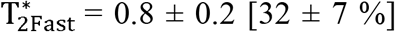, 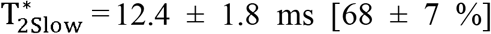).^15^ The average muscle absolute concentration (after T_2_^*^ correction) was 28.6 ± 3.3 mM, which is also in agreement with reported biopsy results.^7^ In terms of T_2_^*^ spatial distribution, the slow fraction is larger and slower in areas with large blood vessel, which makes sense. In blood vessels, the motion is less restricted compared to muscles and spins move more freely that resulting in longer T_2_^*^ values and the time averaged QI is minimal, which means that the fast component fraction is also minimal.

One can conclude at least two important reasons for performing a relaxation study. First, relaxation corrected data provides a better estimation of absolute concentration, as shown by better agreement with biopsy results (see the preceding paragraph). In Figure 7, it has been shown that concentration was increased after the correction. Additionally, one can see that even with UTE measurement, a relaxation bias might be present due to the difference in decay mode within the reference bottles. The second use of T_2_ mapping is the extra information that can be utilized to assess potential physiological changes. Nevertheless, one needs to be aware that voxel-wise T_2_^*^ correction may also result in overestimating ^23^Na concentrations in limited regions, where a hard to avoid partial volume effect exists. This was seen in this study within areas where the main blood vessels start bifurcating to smaller vessels (see the regions of very large difference in Figure 7C).

According to the repeatability analysis, the proposed method showed high repeatability (CV = 4%, ICC = 0.98). The repeatability CVs and ICCs after T_2_^*^ correction were close to the 3D MRI results (CV = 5%, ICC = 0.95 vs CV = 6% and ICC = 0.89). Thus, the proposed MRSI might be a potential method to estimate the T_2_^*^ values and the absolute sodium concentration (free from T_2_^*^ bias) within a reasonable scan time. Since ^23^Na signal suffers from low SNR, with longer data acquisition, scanning with large voxels or at high fields is performed to maintain good SNR within a reasonable scan time. Compared to other techniques at 3T, the proposed MRSI method acquires 64 data points with an acceptable spatial resolution (2.5 mm^2^, nominal in-plane) and acquisition time (15 minutes and 24 seconds). Since the FID data acquired at different time points with different phases, ΔB_0_ correction is also feasible with this method without extra scans. To ensure a proper biexponential fitting for T_2_^*^ estimation, the prolonged acquisition sampling duration allows for acquisition of the entire FID. This is important for areas of longer T_2_^*^ values like reference phantoms.

A limitation of this method is its large dependency on the T_1_ value of the scanned tissues. In order for the OVS to perform correctly, the sequence needed to be applied with TI = ln(2) x tissue’s T_1_ (close approximation when TR>>T_1_). In this study, we used a TI of 20 ms because muscles have a ^23^Na T_1_ of 29 ms, as previously measured at 3T.^4^ To suppress tissues with shorter T_1_, such as cartilages, the TI has to be reduced, which may be technically challenging. However, one can make sure that no such tissues are within the active region of the receiving coil. For instance, we avoided getting residual signals from the knee by keeping it outside the ^23^Na-coil field. Here, we evaluated the technique in healthy muscles. However, T_1_ might change with diseases. Although it is expected that this OVS technique would still achieve good suppression even with slightly T_1_ deviations, future studies might be needed to confirm this. In this study, no ΔB_1_ correction was implemented. However, the coil B_1_ mapping was performed using a GRE sequence with the double angel method.^32^ Since the ΔB_1_ map was very homogeneous, no ΔB_1_ correction was conducted. The ^23^Na-coil ΔB_1_ map can be found in Supporting Figure S1. Although the sample size may be a limitation in this study, there is no reason to anticipate a large difference in our results with larger sample size.

For this study, we used a volume coil placed along the z-axis of the scanner. Thus, the spatial B_1_ variation is expected to increase along the z-direction, where the OVS bands are applied. Together with inversion pulses, OVS bands make the reduction of the longitudinal magnetization depends mainly on the T_1_ and is independent of spatial B_1_ variations. Additionally, the broad-BW OVS pulses were applied with a B_1_ peak of 29 μT, 2.56 ms duration, and a thickness of 100 mm that resulted in no SAR issues.

The considerable regional variability within T_2_^*^, their fraction maps, and the increased accuracy of concentration maps demonstrate the potential for future characterization of ^23^Na in conditions such as muscle diseases, diabetes, cancers, strokes, cartilage degeneration, and to evaluate physiological interventions. The reduction in scan time will increase the technique’s availability and reduce motion artifacts. Technically, while still using a long enough TR (4-5 times T_1_), a shorter TR than that used in this study can be used, which will accelerate the acquisition further. In addition to ensuring a full T_1_ recovery and maintaining a low SAR, we used a very long TR because we wanted to study the existence of any long decaying species. However, no such decay has been noticed within the scanned area. Thus, we would suggest using a TR of about half of the value used here. Additionally, scanning with shorter TEs, and with higher sampling frequency could enhance the fitting quality further. Beyond ^23^Na, the proposed MRSI technique might be useful for ^1^H applications such as imaging dental tissues, which characterized by a very fast T_2_^*^. Measurements of other fast decaying nuclei such as ^13^C, ^31^P, or ^15^N may benefit from the proposed technique as well.

## CONCLUSIONS

The proposed method allows fast data collection for measuring sodium concentrations and T_2_^*^ values at high reliability. This may facilitate evaluating pathological and physiological changes related to the ^23^Na concentration and T_2_^*^ values within a clinically feasible time, and with a good spatial resolution at 3T.

## ACKNOWLEDGMENTS

The study was supported by the Indiana CTSI and funded in part by grant #UL1TR001108 from the NIH, NCATS, CTS Award, as well as a pilot grant by the College of Health and Human Sciences, Purdue University.

## SUPPORTING INFORMATION

Additional supporting information may be found online in the Supporting Information section at the end of this article.

**FIGURE S1** Sodium coil B1 map. The map was generated using a ^23^Na-GRE sequence and the double angle method. A large phantom (15 cm diameter, 300 mM) was scanned with TE/TR: 1.9 ms/120 ms, FA: 45°/90°, 224 averages, resolution: 3 x 3 x 30 mm^3^, FOV: 192 x 192 mm^2^. The normalized B1 map is very homogeneous. Thus, no B1 correction was performed

