## Supporting Figure S1 for "Fast in vivo ^23^Na imaging and T_2_^*^ mapping using accelerated 2D-FID magnetic resonance spectroscopic imaging at 3 T: Proof of concept and reliability study"

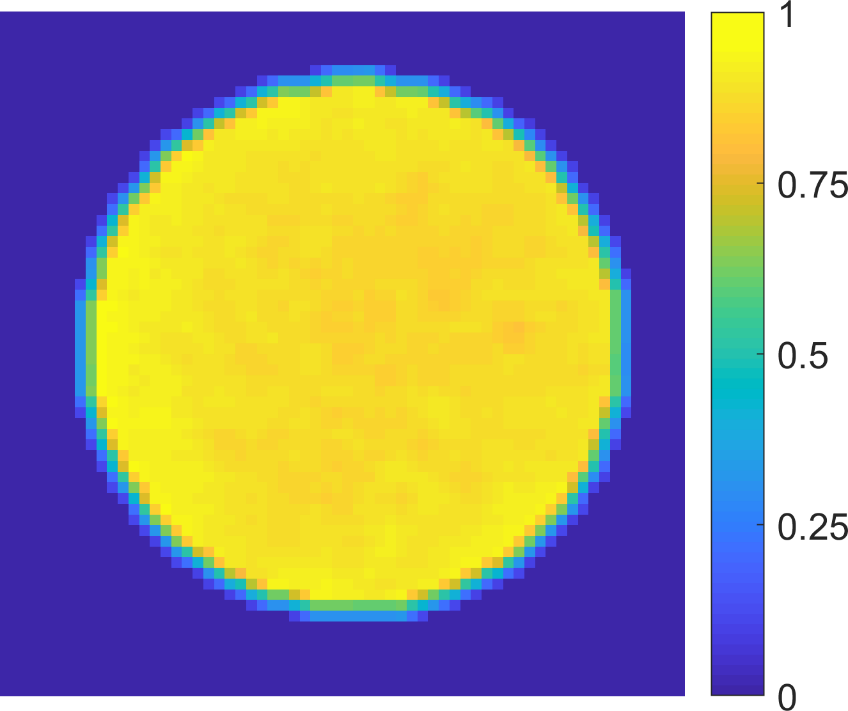

**FIGURE S1** Sodium coil B_1_ map. The map was generated using a ^23^Na-GRE sequence and the double angle method. A large phantom (15 cm diameter, 300 mM) was scanned with TE/TR: 1.9 ms/120 ms, FA: 45°/90°, 224 averages, resolution: 3 x 3 x 30 mm^3^, FOV: 192 x 192 mm^2^. The normalized B_1_ map is very homogeneous. Thus, no B_1_ correction was performed
